# Large Numbers of New Human Paralogs Discovered

**DOI:** 10.1101/2025.10.15.680306

**Authors:** BK Pradeep, Weixia Deng, Mahsa Askary Hemmat, Issak Daniels, Robert L. Jernigan

## Abstract

The identification of paralogs is critical for understanding protein evolution, function, and for drug design, yet many human proteins remain unannotated and poorly classified. Sequence-based homology detection alone often fails to detect distant paralogs, especially in the “twilight zone” or beyond, regarding sequence identity. Here we present an integrated homolog detection framework that combines results from BLASTp, MMseqs2, Foldseek, and the large protein language model-based tool PROST, followed by validation based on comparison of structures and for enzymes comparison of the specific structures of the catalytic residues. Using all-versus-all exhaustive comparisons across the 20,647 human proteins, we systematically identify novel paralogs and assess their catalytic residues for two serine protease clans. We discovered 14 previously uncharacterized human serine carboxypeptidases, validated against experimentally determined PDB structures, with 11 of these displaying conserved catalytic triads. We further identify 203 new paralogs for human kinases, with 163 of these in the major clusters that represent previously uncharacterized kinase subtypes and 30 putative novel human transcription factors. Across both serine protease subtypes, structural alignments enable the prediction of the previously unknown catalytic residues for those lacking UniProt annotations of active site residues. By integrating sequence, structure, and LPLM embedding-based approaches, the framework enables the discovery of surprisingly large numbers of unknown paralogs, permitting defining catalytic residues, and expands the understanding of protein functional landscapes. These findings provide the foundation for a large number of future functional, evolutionary, and therapeutic investigations.

## Introduction

Finding the relationships among different genes/proteins – particularly identifying paralogs (genes derived from duplication events within a single genome) - is fundamental for understanding biological function. The ability to do this relies critically on reliable homolog identification methods and the availability of all sequences. Advances in sequencing technologies have driven down sequencing costs, leading to an unprecedented explosion in sequence data.^1,2^ Recently, breakthroughs in AI-driven deep learning protein structure models, such as AlphaFold.^3,4^ RoseTTAFold,^5^ DeepFold,^6^ OmegaFold,^7^ as well as protein language models such as ESMFold,^8^ and ProteinBERT^9^ have generated large-scale structure predictions with accuracies comparable to experimentally determined structures.^10,11^ Despite these, functional annotations remain incomplete for a significant fraction of proteins.^12,13^ Moreover, there are huge inconsistencies and other differences in the annotations in different databases. These may originate from various experimental results, from inconsistencies in traditional similarity-based function predictions, phylogenomic anomalies, and even from sequencing artifacts.^14^ Compounding this challenge, many proteins, even in some closely related genomes, have not yet been annotated. Yet, the convergence of the growing sequence data and new homolog detection methods offers the unprecedented opportunity to close this gap by producing new inferences for the functional relationships among proteins. In addition, the newer reliable structure predictions provide a direct way to validate the predicted sequence relationships. Learning about all paralogs is important in designing drugs because if not all paralogs were used in testing a drug, it may not be fully effective against all desired protein targets, or a new drug might interact with some of the paralogs in unexpected ways, causing unexpected side effects.

Traditional sequence-based homolog detection relies on computing a significantly similar alignment score to infer the reliability of transferring annotations from the known protein to the target. However, because protein structures are more conserved than sequences, many homologs are likely to have sequence identities well below the range of reliable detections. into and below the usual twilight zone boundary of 25-30% sequence identity.^15,16^ However, proteins found in this lower range of sequence identity can be either similar or completely different, thus complicating homolog detection. There are now significantly improved new homolog detection methods, which can, however, reliably discover new homologs and overcome many of the previous problems. Recently, homolog detection tools powered by the Large Protein Language Models (LPLMs) show a remarkable promise in finding new homologs, because they have digested huge numbers of sequences to capture overall the biophysical, biochemical, evolutionary, phylogenetic, structural, and mutational insights within their embeddings.^17^ For simpler organisms like *Bacillus subtillis*, one of these methods PROST^15^ enables a nearly complete set of functional annotations; see https://bit.ly/prost-bsubtilis (slow loading). In higher organisms like humans, it appears, however, that the challenge requires an integration of multiple approaches. There are some existing databases of paralogs: ENSEMBL,^18^ Paralog Explorer,^19^ and FlyBase^20^ are some of the noteworthy ones. The methods used to develop these databases were only sequence similarity and phylogenetics.

Here, we introduce a novel integrated framework that enables the discovery of significant numbers of new human paralogs based on information from sequence-structure relationships, protein language models, and functions. Specifically, we use BLASTp.^21^ MMseqs2,^1^ Foldseek,^22^ and PROST^15^ in this integrated method for homology identification. BLAST and MMseqs2 are sequence-based homolog detection tools; whereas, Foldseek is a structure-based tool. PROST leverages the embeddings generated by the large protein language model ESM1b.^23^ Using this integrated approach, we perform an all-vs-all search for the paralogs of all human proteins. For the sequences themselves, we use the UniprotKB^24^ reference proteome database to obtain a non-redundant set of 20,647 human protein sequences. Of these, 20,208 proteins have predicted structures available in the AlphaFold Database,^4,25^ enabling structural comparisons for almost all pairs, which we use for validating the new paralogs. As examples of the new paralogs we show results for proteases, kinases, and transcription factors. Their structures are clustered based on the structural similarities using TM-scores from TM-align.^26^ We chose the proteases because they have been so intensively studied and the kinases and transcription factors because of the relatively larger numbers of these. The validation compares structures within a cluster with the structures of the known members of the cluster. To ensure similar functions of the proteases, we investigate the corresponding positions of the active site amino acid triads. For some cases this enables new identifications of the active site residues not only for the new paralogs but also for some of the previously identified proteases with prior annotations of their active sites. These structural comparisons are carried out using both AlphaFold2 predicted structures as well as experimental crystal structures.

## Results

### 17 Newly Discovered Human Serine Carboxypeptidases

MEROPS^27^ is a specialized database for proteases that classifies human proteases into 10 different subtypes; Hedstorm^28^ further categorized serine proteases into clans. For our analyses, we use the previously known 108 chymotrypsin-like serine proteases (with catalytic triads H-D-S) and 13 serine carboxypeptidases (with S-D-H triads) as the reference set. The integrated homolog searches and structural validations using bi-directional TM-scores ≥ 0.5 yield 132 paralogs, which are clustered into four major groups shown in supplemental **Fig. S1** (see details in supplemental Table S1). The cluster with 14 of the 17 newly discovered paralogs is shown in **Fig. 1**. The novel serine carboxypeptidases from **Fig. 1**, are Q8WTS1, Q5VYY2, Q8TB40, P07098, P07099, Q5VXJ0, P38571, Q5W064, Q5VXI9, Q5EB52, Q92597, Q8IUS5, Q9H6B9, and Q9BV23. Of these, five proteins - P07098, Q5VXJ0, Q5W064, Q5VYY2, and Q5VXI9- are not listed in the MEROPS^27^ (database that records the proteases across various organisms). Two prominent clusters (see supplemental **Fig. S1**, blue) comprise canonical chymotrypsin-like serine proteases, exhibiting the conserved catalytic triad of His-Asp-Ser. Notably, there are several proteins in these clusters with missing active site annotations. We also predict what these residues are by using a multiple structure alignment approach described in the methods. Two clusters in the supplemental **Fig. S1** (green), are **serine carboxypeptidases**, exhibiting the conserved catalytic triad of **Ser-Asp-His**. The modular classification of these novel discoveries in the cluster suggests that they have a similar canonical catalytic triad for most as predicted and shown in **Table 1A**. This distinct cluster of serine carboxypeptidases has no overlap with other clusters, indicating an expansion of the number of functional subclasses. Furthermore, the conserved structures of serine carboxypeptidases, as shown in supplemental **Figs. S2C**, and **S2D**, provides evidence for two different structural types, both having the same function.

**Figure 1.**
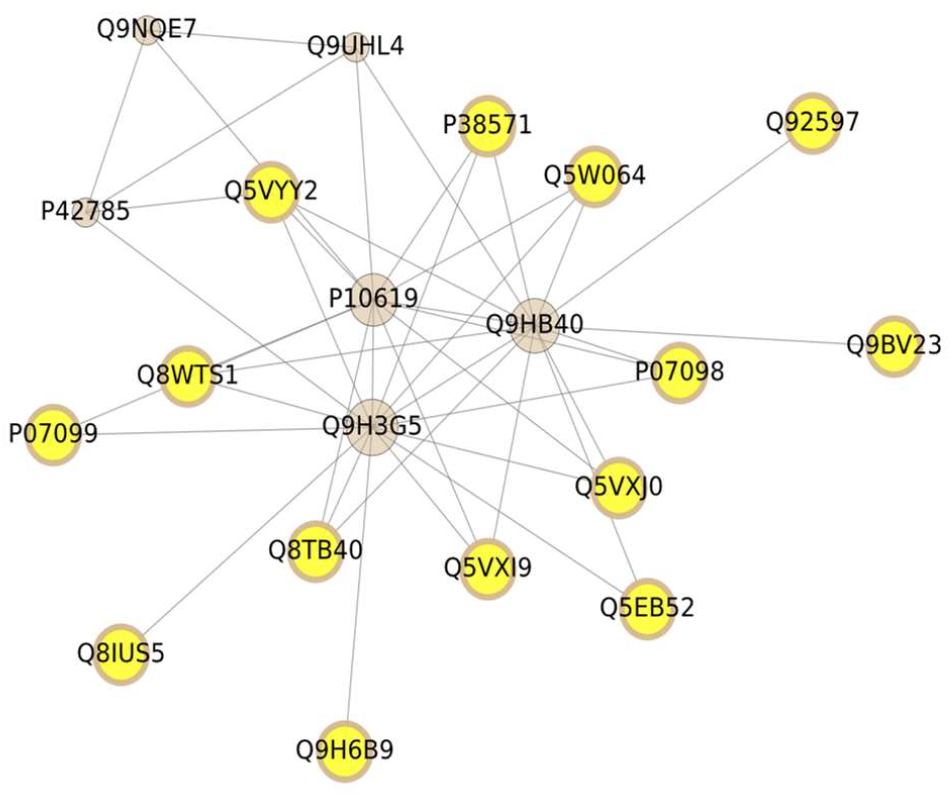
Network representation of 20 human serine carboxypeptidase structures (AlphaFold generated) using the Louvain community detection algorithm.^40^. The clustering is based on structure similarity with both bidirectional TM-scores being ≥ 0.5. Solid gray nodes are previously known serine carboxypeptidases having canonical catalytic triads of His-Asp-Ser, whereas those highlighted in yellow are newly discovered proteases. Node sizes are proportional to the number of neighbors connected by edges, except for the newly identified ones, all with similar node sizes. Structural alignment s of these 14 new proteins with the 3 central known proteases using PyMOL^41^ reveals high fold conservation with an average whole structure RMSD of 4.91 Å ± 1.01 Å (see supplemental **Fig. S2D**). Sequence similarity for these 17 proteases computed by Clustal Omega^41^ pairwise alignments using the BLOSUM62 matrix is strong showing 51.77 % ± 7.60 % (see **Fig. S3**).

**Table 1.** Active site residue predictions for previously unannotated serine proteases. All now have predicted active site residues. Orange highlighted rows are predictions with weak structure validation (large RMSDs), and blue highlighted rows are predictions for previously unannotated serine proteases in the two separate large blue clusters in **Fig. 2**. Additional information gleaned from literature searches is given in Table S5.

|  |  |  |  |  |
| --- | --- | --- | --- | --- |
| A. Active site residue predictions for novel serine carboxypeptidases compared with known protein Q9H3G5 |  |  |  |  |
| UniProt ID | Name | UniProt active site residues | Predicted active site residues | Catalytic Triad RMSD (Å) |
| P38571 | Lysosomal acid lipase/cholesterol ester hydrolase | S174, H374 | S174, D345, H374 | 0.27 |
| Q8TB40 | (Lyso)-N-acylphosphatidylethanolamine lipase | None | S146, D290, H320 | 1.70 |
| Q5EB52 | Mesoderm-specific transcript homolog protein | None | S145, D284, H314 | 2.50 |
| Q8WTS1 | 1-acylglycerol-3-phosphate O-acyltransferase ABHD5, experimentally shown to be a pretease<sup>44</sup> | None | S163, D301, H327 | 6.11 |
| P07099 | Epoxide hydrolase 1 | D226, Y374, H431 | S229, E404, H431 | 1.63 |
| Q5VXJ0 | Lipase member K | S172, D343, H372 | S172, D343, H372 | 0.19 |
| P07098 | Gastric triacylglycerol lipase | S172, D343, H372 | S172, D343, H372 | 0.12 |
| Q5W064 | Lipase member J | S141, D312, H341 | S141, D312, H341 | 0.15 |
| Q5VYY2 | Lipase member M | S186, D357, H386 | S186, D357, H386 | 0.21 |
| Q9BV23 | Monoacylglycerol lipase ABHD6 | S148, D278, H306 | S148, D278, H306 | 0.24 |
| Q5VXI9 | Lipase member N | S173, D344, H373 | S173, D344, H373 | 0.21 |
| Q8IUS5 | Epoxide hydrolase 4 | D169, Y281, H336 | S161, D307, H336 | 11.0 |
| Q9H6B9 | Epoxide hydrolase 3 | D173, Y281, H337 | S165, D307, H337 | 11.2 |
| Q92597 | Protein NDRG1 | None | S133, D261, H194 | 12.9 |
| B. Active site residue predictions for chymotrypsin-like serine proteases against references P05981 (light-blue), and P00746 (blue) |  |  |  |  |
| P00738 | Haptoglobin | None | H208, D246, S357 | 5.80 |
| P22891 | Vitamin K-dependent protein Z | None | H223, D261, S355 | 4.98 |
| Q6PEW0 | Inactive serine protease 54 | None | H66, D122, S223 | 6.88 |
| A4D1T9 | Probable inactive serine protease 37 | None | H55, D99, S185 | 9.31 |
| P00739 | Haptoglobin-related protein | None | H150, D188, S299 | 5.56 |
| P20160 | Azurocidin | None | H58, D115, S216 | 6.72 |
| A0A1B0GVH4 | Serine protease-like protein 51 | None | H79, S193 | 5.20 |

### Multiple Structural Alignment of Newly Discovered Proteases with a Previously Annotated Protease Identifies Missing or Alternative Catalytic Residues

To examine catalytic site conservation, we perform multiple structural alignments for the 20 proteins in the cluster (**Fig. 1**) using FoldMason, with the central known human serine carboxypeptidase, **Q9H3G5**, as the structure reference. By anchoring the alignments to the canonical catalytic triad residues (Ser-Asp-His) of Q9H3G5, we systematically inspect the corresponding residues for all 14 novel discoveries (**Table 1A**). We do the same to assign the missing catalytic residues for 7 of the previously known chymotrypsin-like serine proteases with missing catalytic residue annotations by using the reference proteins **P05981** (light-blue cluster) and **P00746** (blue cluster) shown in supplemental **Fig. S1**.

This reveals that four novel carboxypeptidases previously lacking annotated active sites (at the top of Table 1A) in UniProt can now be assigned the SDH catalytic triad. The correct assignments of sequence positions for previously known active site residues are also obtained and shown (see columns 2 and 3, Table 1), which further validates the functional assignments in a highly specific way. Also, for **P38571**, which previously had only two catalytic residues annotated, it now has a completed catalytic triad. **Figure 2** shows the overlapping conserved catalytic residues highlighted in blue for reference active site structures, and yellow for predicted active site residues having good RMSDs, as low as 0.12 Å. Apparently, **P07099** has glutamic acid corresponding to the reference aspartic acid, showing this alternative active site residue. Despite being annotated to have D-Y-H as catalytic residues in UniProt, our method suggests S-E-H based on these residues’ small RMSD from the reference serine protease triad. By contrast, putative carboxypeptidases **Q8IUS5, Q9H6B9**, and **Q92597** do not show Ser-Asp-His near the reference triad, suggesting either possibly an alternative catalytic residue or a false homolog; however, their structural similarity to experimentally verified proteases strongly suggests their possible characterization as another protease subtype.

**Figure 2.**
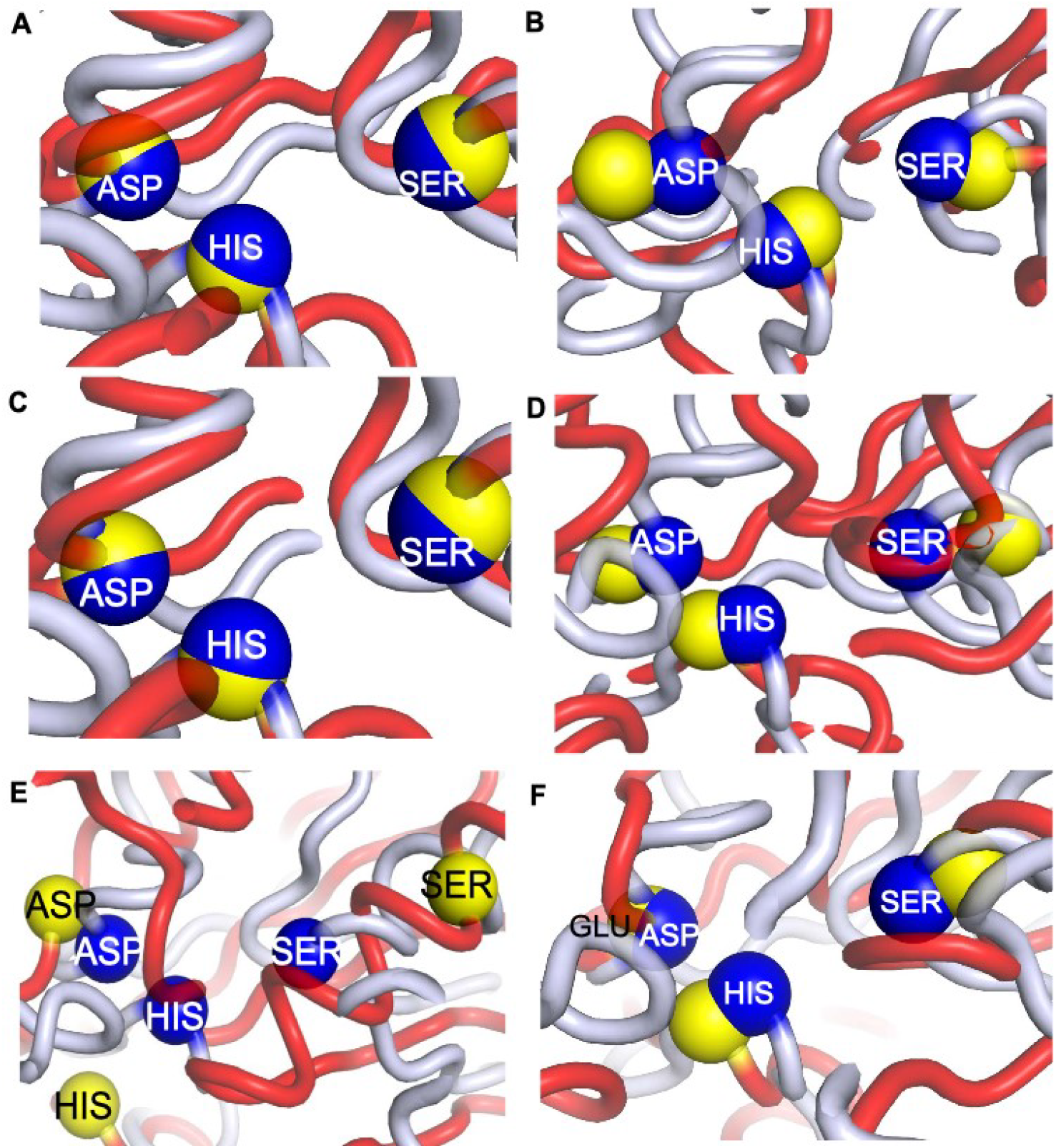
Structural alignments of the active site triads of the known and newly identified human serine carboxypeptidases. Pairwise structural alignments of putative novel serine carboxypeptidases A. **P07098**, B. **Q8TB40**, C. **P38571**, D. **Q5EB52**, E. **Q8WTS1, and** F. **P07099** with the known serine protease (**Q9H3G5** aligned using PyMOL for the triads. The three known catalytic triad residues (Ser-Asp-His) in Q9H3G5 are shown as blue spheres. Predicted catalytic residues in the novel paralogs, inferred via multiple structure alignment using Foldmason^39^, are shown as yellow spheres. The RMSD values of **0.12 Å, 1.70 Å, 0.27 Å, 2.50 Å, 6.11 Å, and 1.63 Å** between reference and the corresponding targets (A-F) highlight the strong structural conservation for the catalytic sites. Only E shows larger differences and it is possible that this could be because of the dynamics of the structure.

Notably, seven of the previously known chymotrypsin-like serine proteases had missing active site residues. We implement the same multiple structural alignment to examine the corresponding residues **(His-Asp-Ser)** in these proteases near the catalytic triads of the reference proteins for **P05981** (light-blue cluster) and **P00746** (blue cluster) shown in supplemental **Fig. S1**, and reported in **Table 1B**. Apparently, the structure alignments of catalytic residues between the reference and the target using PyMOL shows structural divergence with somewhat larger RMSDs but still enables identifying the active site triads.

### Validation of New Carboxypeptidases

We identify 14 previously unrecognized human serine carboxypeptidases using our integrated homolog detection pipeline and structural validation using functional clustering based on TM-scores with the Louvain community clustering algorithm (see **Methods**). Because AlphaFold structures are predicted structures, we have also compared each candidate directly with experimentally determined PDB^29^ structures, and 10 out of the 14 novel serine carboxypeptidases have structural similarities to experimental proteases. This additional step ensures that the observed similarities are not prediction artifacts but are grounded in independent experimental data. Four of these novel proteins match structures annotated as hydrolases or a peroxidase or lipase-like enzymes (see **Table 2**, bottom four rows) with good TM-scores, revealing perhaps either an unexpected similarity between serine proteases and these other enzymes or false identifications. The observed relationship between carboxypeptidases and lipases arises with the shared α/β-hydrolase fold but with different catalytic functions.^30, 31^

**Table 2.**
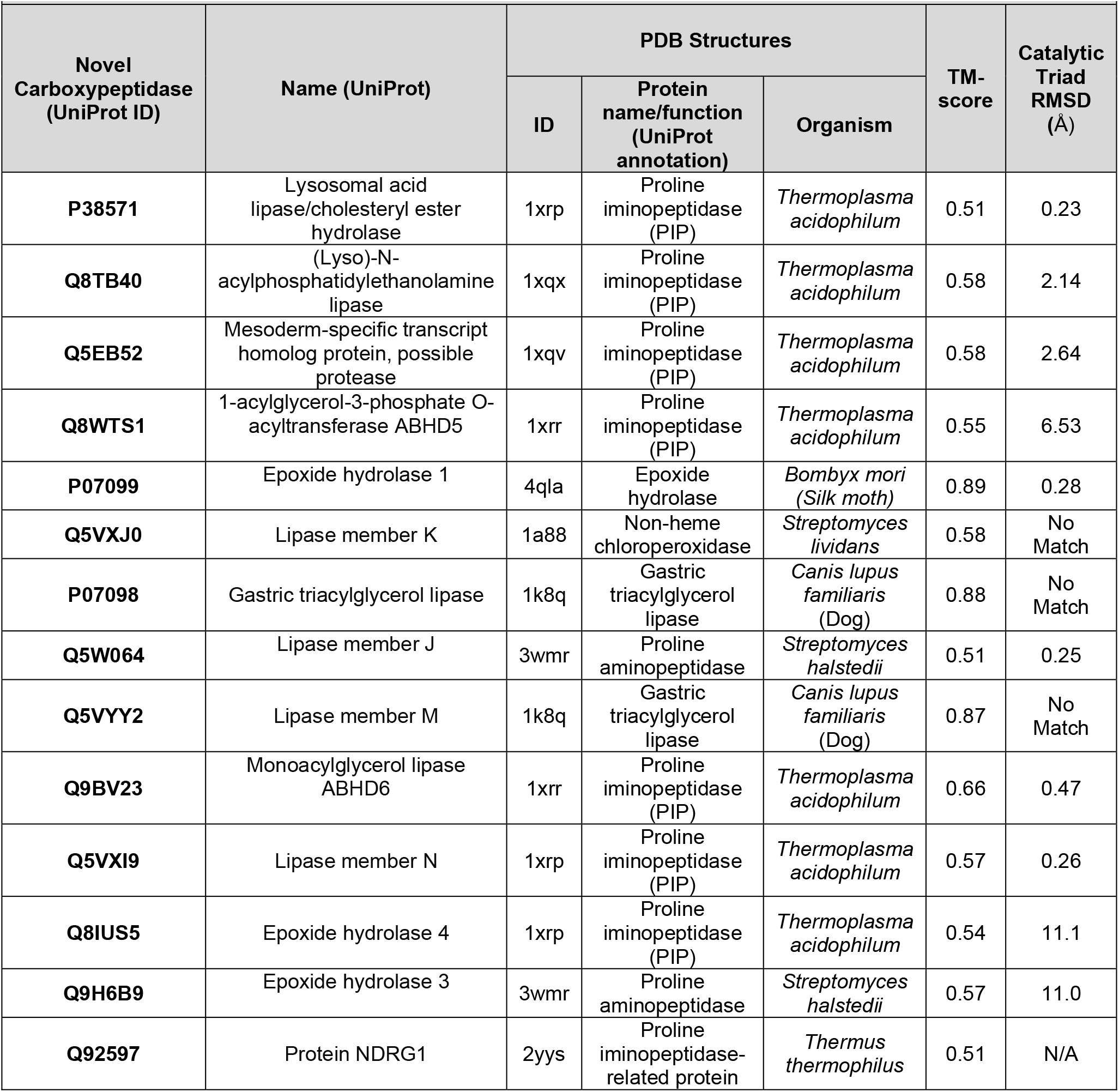
Newly Discovered Serine Carboxypeptidases, with Experimentally determined PDB structural homologs.

These novel paralogs are further validated by the conservation of their catalytic residues, as shown by representative structural alignments, based on the catalytic residues, blue for the pdb reference structures and yellow for the new paralogs, shown in **Fig. 3A-C**. The alignments have RMSDs as low as 0.23 Å. Furthermore, we also predict active site residues for a total of 21 proteases (14 novel serine carboxypeptidases, 7 active-site-unannotated chymotrypsin-like serine proteases) (see **Table 1**). Notably, proteins with high or missing RMSD values in Table 2 (Q8IUS5, Q9H6B9, and Q92597) also show high RMSD values in Table 1, adding additional support for their being different. However, no catalytic triad matches are found for the protein pairs Q5VXJ0–1a88, P07098–1k8q, and Q5VYY2–1k8q suggesting that, despite being annotated as lipases or peroxidase in Table 2, these proteins should be classified as serine carboxypeptidases, supported by their catalytic triad agreement with known proteases (**Table 1**). The catalytical triads are not quite the same. The validation with the experimental structures confirms the new paralog annotations but also further validates this identification by aligning the correct active site residues with the reference experimental structure.

**Figure 3.**
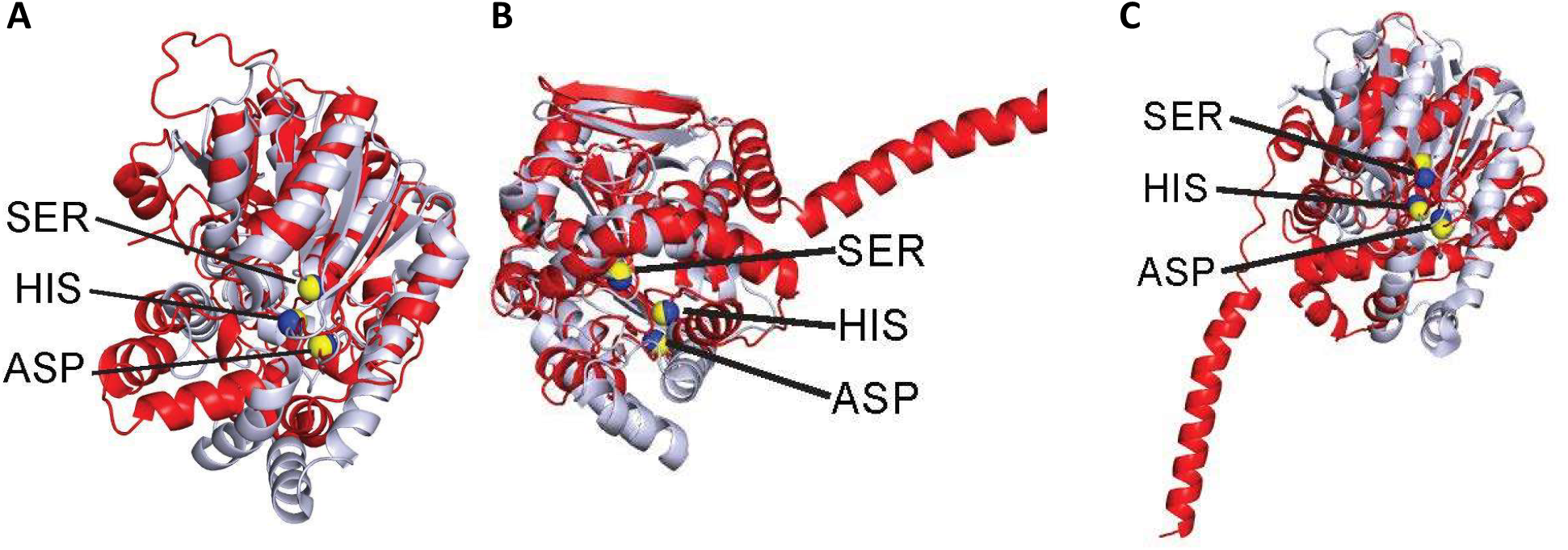
Structural alignment of AlphaFold2 predicted novel serine carboxypeptidase structures with homolog PDB structures using PyMOL^42^ of the triads. All 14 newly discovered human serine carboxypeptidases have significant structural similarities to their experimental homologs (TM-scores ≥ 0.5 bidirectionally; see entries in Table 1). 10 of the 14 new discoveries exhibit a clear conservation of the catalytic Ser-Asp-His triad, which aligns closely with the active site residues of their experimental homologs. **A**. Protein pair Q5W064-3wmr (triad RMSD = 0.25 Å), **B**. Protein pair Q9BV23-1xrr (RMSD 0.47 Å), and **C**. Protein pair Q5EB52-1xqv (RMSD 2.64 Å). These novel proteins showing significant structural similarities to experimentally determined proteases ensure that these are indeed serine proteases.

### Application to Human Kinases Yield Large Numbers of New Paralogs

This integrated homolog detection to the 580 annotated human kinases (QuickGO), yields 5,048 novel paralog proteins in 78,757 pairs. After structural validation (TM-score ≥ 0.5 bidirectionally), the remaining 582 unique proteins with these good TM-scores show 417 of these clustering into 12 groups, shown in supplemental **Fig. S8**. This implies a high level of diversity within the family.

This results in 203 newly discovered kinases. Of these, 163 are retained in the main larger clusters shown in **Fig. 4**. All these putative novel kinases in the various clusters are listed in Supplemental **Tables S2** and **S6**. Strikingly, clusters with large numbers of yellow-highlighted paralogs in **Fig. 4** represent novel kinases that were not previously annotated, suggesting overall a substantial expansion in the extent of kinase-mediated regulation.

**Figure 4.**
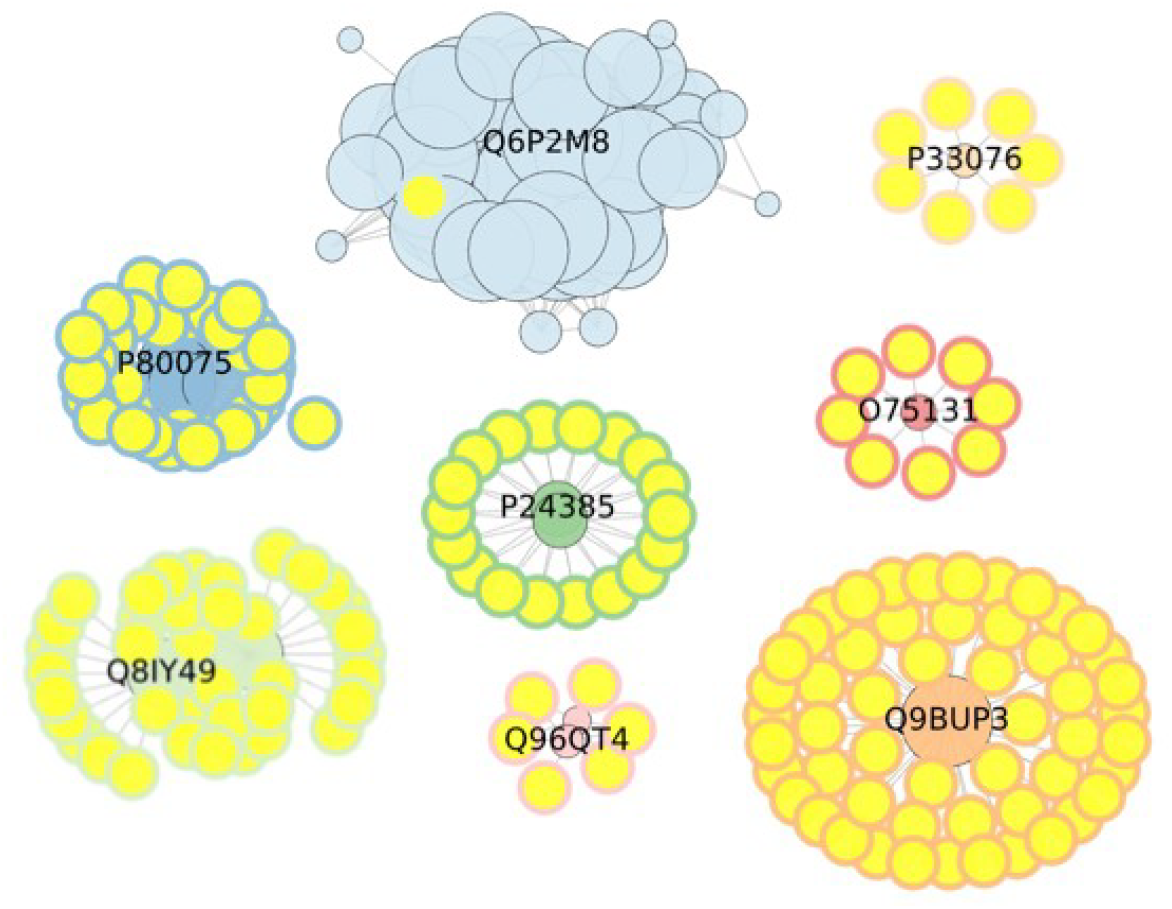
Eight clusters of 417 structurally similar (TM-score ≥ 0.5 in both directions) human kinase paralogs showing their complex relationships. 582 proteins with TM-scores ≥ 0.5 bidirectionally are clustered by using the Louvain community detection algorithm, and clusters larger than 7 are shown. 163 newly identified kinases are highlighted in yellow with different outline colors are shown together with their central identified known kinase.

### Application to Transcription Factors

In an additional example, we investigate human transcription factors for 1,373 reference proteins identified in PAN-GO^32^. We identify 5,061 additional paralogs from our integrated homology search. With structural validation and removing clusters having fewer than eight members, the remaining 417 putative paralogs fall into 9 clusters, as shown in supplemental **Fig. S9**. There, 30 novel transcription factors are identified and shown yellow-highlighted (see supplemental **Tables S3** and **S7**). Initial investigation shows the difficulty in validating these structurally because of their common modularity, with different homologs having different numbers of modular structure components.

Our integrated homology search and structural clustering approach provides a framework for uncovering candidate subfamilies for any class of proteins, guiding future biochemical and functional validation efforts, providing a clear way to expand the landscape of paralogs.

## Conclusions and Discussion

In this study, we have presented an integrated homolog detection framework that combines sequence-based BLASTp and MMseqs2 with structure-based Foldseek, and LPLM-based PROST approaches with downstream structural and active site validations. Applying this strategy to the human proteome reveals an extremely large number of previously unrecognized paralogs. In the present limited study, we found 14 novel serine carboxypeptidases, 203 novel kinases, and 38 novel transcription factor candidates. Many of the novel serine carboxypeptidases display strong structural similarities to experimental PDB entries as well as AlphaFold predicted structures. By leveraging multiple structural alignments, we are able to predict catalytic residues for the novel proteases and for some known proteases that were missing catalytic residue annotations, thereby filling critical gaps in current databases. There are some inconsistencies in the functions previously reported compared to the functions found here. This will be further investigated with clustering based on a variety of other scorings in addition to the scoring here by structure comparisons.

Our results present several different insights. First, there is the observation of the structural diversities defining protein family subtypes, even within well-studied families such as serine proteases and kinases. This raises a question about the possible differences in their dynamics, localizations, and specificities, vis a vis their functional similarities. Second, the emergence of numerous separate kinase-like clusters demonstrates that structural and functional space is broad yet more interconnected than may have been previously presumed. Importantly, since analyses underscore the value of combining multiple orthogonal homolog detection methods as different methods perform best for different classes of proteins as shown in **Table 3**. Here, PROST along with Foldseek and BLAST perform better to find novel homologs for the transcription factors, whereas Foldseek performs best for the kinases. Likewise, PROST outperforms others for carboxypeptidases. While sequence similarity remains a cornerstone, the use of the large protein language model embeddings demonstrate large gains. In addition, we have shown how structural and functional residue conservation provide crucial ways to validate sequence-based discovery. It is also surprising that new paralogs are still found with BLASTp.

**Table 3.** Distribution of Novel Paralogs Identified by Different Methods.

| **Human Protein Family** | **# novel paralogs** | **# paralogs pairs (reference, novel)** | **PROST** | **Foldseek** | **MMseqs2** | **BLASTp** |
| --- | --- | --- | --- | --- | --- | --- |
| Carboxypeptidases | 17 | 42 | 41 | 33 | 9 | 8 |
| Kinases | 203 | 422 | 285 | 358 | 190 | 80 |
| Transcription Factors | 38 | 162 | 133 | 132 | 85 | 132 |

However, functional discrimination of paralogs based on catalytic or binding motifs can be difficult when substrate preferences have different degrees of specificity, or are similar across unrelated enzymes, or lack unique sequence features, as seen in human kinases.^33^ In contrasting, there are structural modules in the case of transcription factors such as zinc finger proteins.^34^ Yet, the functional similarity is not solely determined by static structures: proteins with related functions often require conservation of their underlying dynamics, as exemplified by RNase enzymes within subfamilies.^35^

While paralog identification is an important approach for protein functional annotation, establishing such relationships among proteins is challenging. So, an integrated approach of comparative genomics, such as evolutionary and functional information and manual curation, is necessary to separate out paralogs from multifunctional families.^36^ Nevertheless, these limitations present opportunities for experimental characterization of these candidates to uncover novel functional subtypes, refine existing classification, and provide insights into protein evolution and drugability. In summary, we foresee that the present framework and its pipeline strengthen protein annotation and opens new possibilities for therapeutic discovery by revealing new relevant candidate drug targets, important for avoiding off-target interactions.

## Methods

### Discovery of Novel Human Paralogs with Integrated Homolog Searches

To systematically identify novel paralogs within the human proteome, we employ an integrated BLASTp-MMseqs2-Foldseek-PROST homolog search method. BLASTp is an extremely well-known local sequence alignment-based homolog detection. MMseqs2 optimizes the exact alignment by first applying consecutive k-mer matches as a pre-filtering step, making it 400 times faster and is more sensitive than BLASTp. Foldseek does structure-based homolog searches by converting the three-dimensional structure information into a 1D-structural alphabet string, followed by MMSeq2 sequence-based search on these strings that represent structures. PROST is our own extremely different method the optimize the use of the LPLM-based ESM1b^23^ sequence embeddings to identify new homologs. Notably this method PROST by itself performed particularly well in the CAFA 5 competition for function predictions, scoring in the top 1.4 percentile.

For each method, we construct a custom database for the 20,647 proteins and perform an exhaustive all-versus-all search to identify the best hits in homolog searches. Then, each method is filtered for the reciprocal hits. Combining the hits from each method and removing the duplicates, the distribution of human paralogs identified by each method for all cases is shown in **Figure 5A**. We use our pipeline to investigate the paralogs in several human protein families – for proteases, kinases, and transcription factors. There are large inconsistencies across databases - for example, 391 peptidases in PAN-GO^31^ and 626 proteases in QuickGO, 448 kinases in PAN-GO and 580 in QuickGO, and 1373 transcription factors in PAN-GO, with only 477 in QuickGO. We use the databases with the larger number of annotations to obtain a reference set for each protein family of interest. We first chose proteases because they are one of the most intensively studied enzyme families, have well-characterized catalytic mechanisms, and critical roles in numerous physiological and pathological processes. We identify 752 annotated proteases from QuickGO^37^ and MEROPS,^27^ of which 618 have annotated active site information in the UniProt database. For the human proteases as our reference set, these methods found an additional 5,463 novel targets with 37,206 unique paralog pairs distributed as shown in **Fig. 5B**. For 580 reference human kinases, the integrated method found 5,628 candidate kinases as shown in **Fig. 5C**. Likewise, using 1,373 transcription factors as reference proteins, 4,236 putative novel transcription factors with 164,166 unique paralogs pairs are identified and are distributed as shown in **Fig**. 5D.

**Figure 5.**
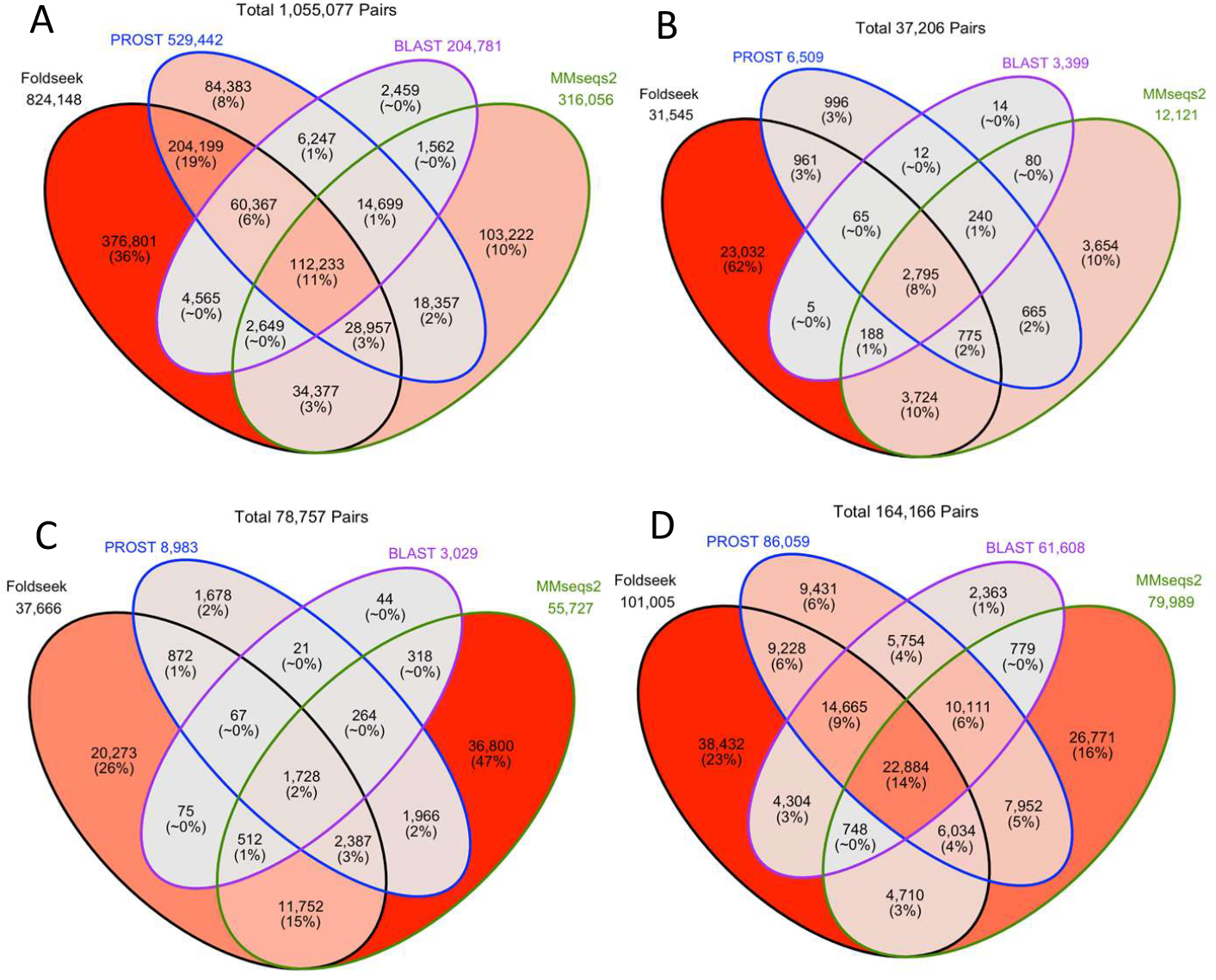
The number of human paralogs found by different methods. The number of paralogs found by different methods, showing total as well as the ones in common for **A. all human proteins, B human proteases, C human kinases, and D transcription factors**. The outline colors of the segments include all the predicted cases for each individual method. These are colored by PROST (blue), Foldseek (black), MMseqs2 (green), and BLASTp (purple). Filtering out the paralog pairs with no reciprocal hits, there are altogether 1,055,077 unique paralog pairs for all human proteins (A), 37,206 unique pairs for 752 reference proteases (UniProt^24^ and MEROPS^27^) and their 5,492 candidate paralogs (B), 78,757 unique pairs for 580 reference kinases and their 5,628 candidate paralogs (C), and (D) 164,166 pairs for 1,373 reference transcription factors and 5,061 candidate paralogs.

### Distribution of Catalytic Residues Among Known Human Proteases

For the known 618 human proteases in the functional annotation of QuickGO,^36^ we retrieve active site residue information from the UniProt database. The resulting distribution (Supplemental **Fig. S10**) reveals that the canonical His-Asp-Ser catalytic triad is one of the most common. This is consistent with the well-established mechanistic paradigm of serine proteolysis. A significant number of proteases, however, lack annotated catalytic residues, suggesting either incomplete curation or functional divergence. Interestingly, beyond the predominant eight kinds of proteases as described in MEROPS,^27^ we observe a range of alternative residue combinations at active sites, indicating some diversity in functional specificity and evolutionary divergence across proteases. Importantly, identifying catalytic residues in the currently unannotated proteases remains a challenge. Computational or structural approaches to infer these residues will enhance our understanding of their biochemical functions and may reveal novel therapeutic targets within this highly versatile enzyme family. For our pipeline to discover novel proteases, we chose two subtypes of serine proteases to investigate: 108 chymotrypsin-like serine proteases with the catalytic triad His-Asp-Ser and 13 serine carboxypeptidases with the catalytic triad Ser-Asp-His as our reference sets.

### Louvain Community Clustering based on TM-Scores

Due to the complex relationships between many proteins in terms of similarity, we separate them into different clusters based on the similarity networks defined by the TM-scores using the Louvain Algorithm.^38^ Here, each protein is represented as a node, and two nodes are connected by an undirected edge if the TM-score is ≥ 0.5 (for both reference and target), forming a network of communities. The clusters with small sizes (< 8 members for proteases, and kinases; < 5 for DNA-binding TFs) were removed to simplify.

One of the ways to detect a community is by modularity of the community, which defines the strength of the network. High modularity means more connections between the nodes within a community but sparse connections between the nodes of different communities. This method maximizes the modularity of the module or the community. The gain in modularity by moving an isolated node *i* into a community *C* is tested. (See supplemental information for more details about the clustering method.

### Structural Alignment using FoldMason^39^

Multiple structural alignment is essential to analyze distantly related proteins. FoldMason is an efficient, highly accurate, and fast method to perform multiple structure alignments on hundreds of thousands of proteins. It leverages the discretized 1D structural alphabets using 3Di interacting residues previously used by Foldseek,^22^ to perform progressive alignment on structures along the guide tree To generate a guide tree, FoldMason first performs pairwise comparison of all input structures using a SIMD-accelerated gapless alignment used in FoldSeek, which is then sorted by their best-to-worst order. The reordered sequences, sorted by their scores, are then aligned using progressive alignment.

### TM-Align and RMSD

TM-align uses both the TM-score rotation matrix and dynamic programming to compute the structural similarities between protein pairs and has been used extensively at CASP for comparing structure predictions.

RMSD compares the protein structures by computing the root-mean-square distances between the corresponding geometric points in the two structures, which have been aligned by optimally rotating and translating one protein relative to the other.^40^

## Source Code

Software is available at: https://github.com/pradeepbkarma/human_paralogs_project

## ACKNOWLEDGMENT

This work was supported by NIH grants R01HG012117, R01GM144961, and R01GM157600.

