## Supplemental Material for "Large Numbers of New Human Paralogs Discovered"

**Supplementary Material**

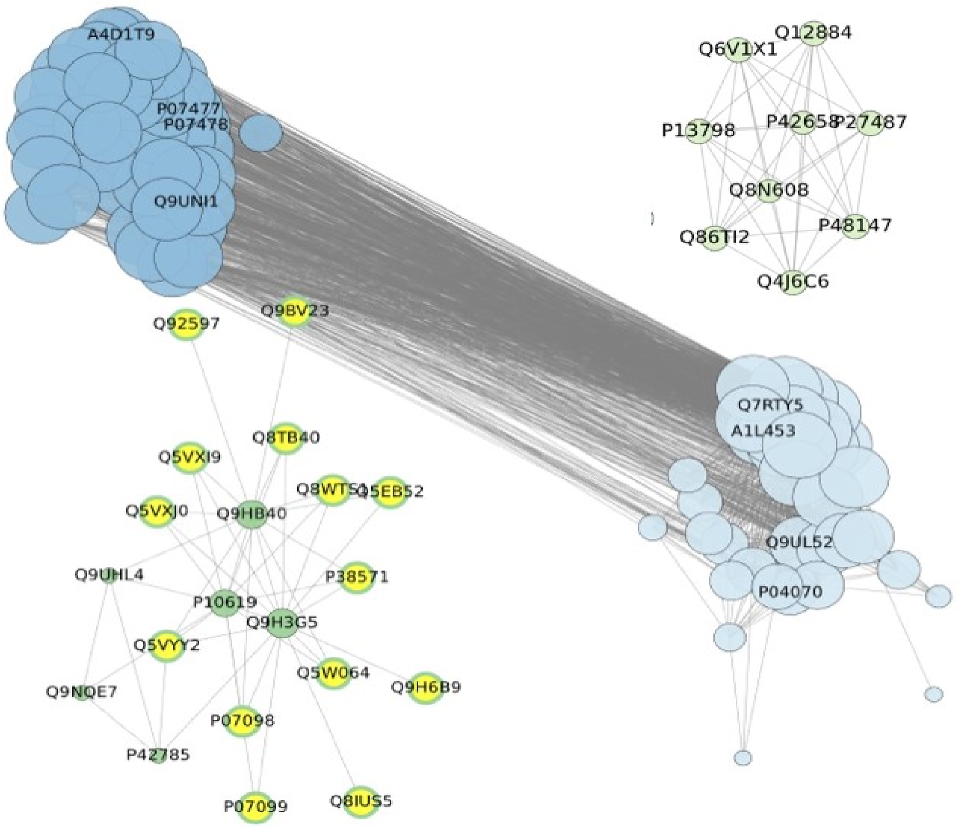


**Figure S1. Modular clusters of human chymotrypsin-like and carboxypeptidase (Yeast-like) serine protease structures with** the members of each cluster having bi-directional TM-scores ≥ 0.5. (All structures used here were generated by AlphaFold2.) Clustering uses the Louvain community detection algorithm (see details below). Proteases with the canonical catalytic triad of His-Asp-Ser are in two clusters (darker and lighter blue); whereas the two with Ser-Asp-His triads are shown in green (lighter and darker). Solid nodes are the previously known proteases, and the yellow filled ones are the newly identified proteases. Node sizes are proportional to the number of neighbors. The dark green cluster on the lower left includes all 14 novel paralogs to the 3 central known serine carboxypeptidases. Structural alignment of these 17 proteases using PyMOL[48] reveals strong structural similarity with a mean whole structure RMSD of 4.91 Å ± 1.01 Å (see supplemental **Figure S2**). Sequence similarity in a multiple sequence alignment using the BLOSUM62 matrix for these 17 proteases is high at 51.77 % ± 7.60 % as shown in supplementary **Figure S3**.

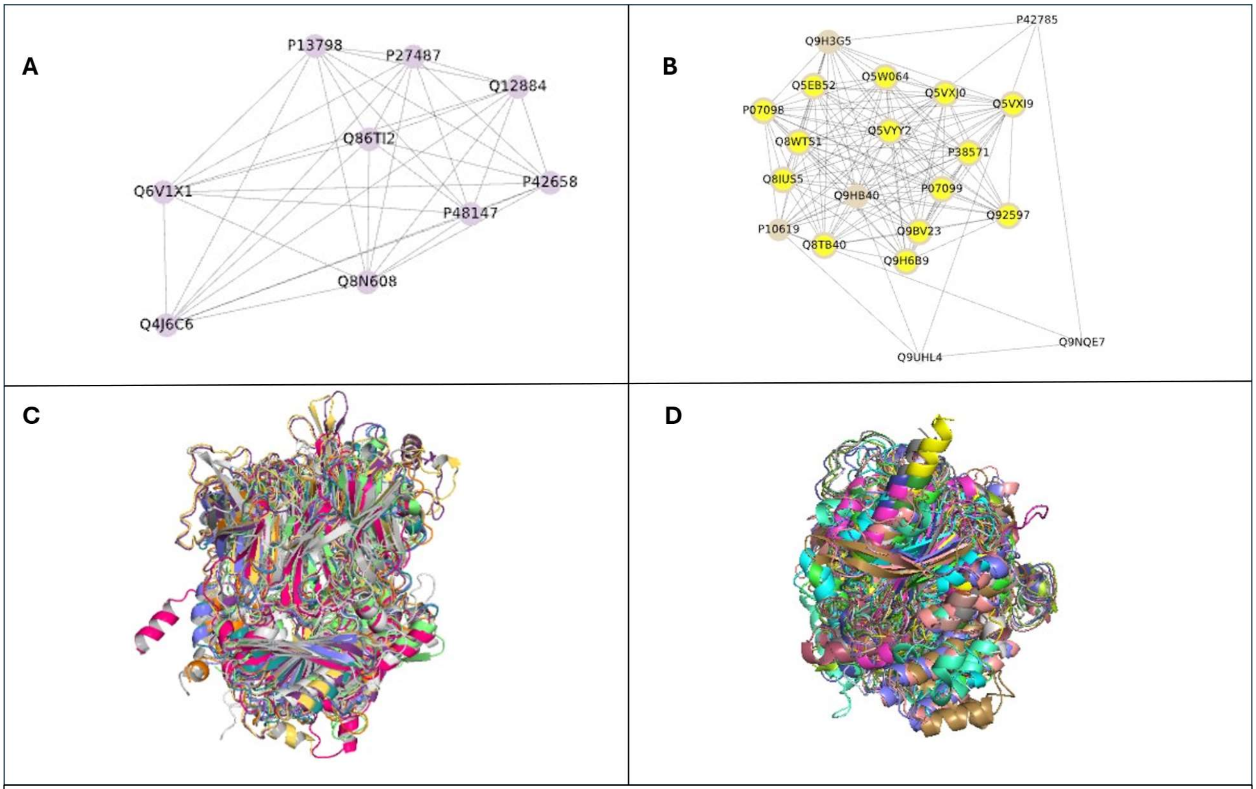


**Figure S2. Structural clustering and alignment of human serine carboxypeptidases. (A, B**) Network representations of 9 (all known types of proteases) and 20 carboxypeptidase Y-like serine protease paralogs, clustered using the Louvain community detection algorithm based on pairwise structural similarity (TM-score ≥ 0.5 for both directions). Solid nodes indicate known proteases; yellow-highlighted nodes represent **putative novel paralogs**. Cluster B contains **14 such novel candidates**. (**C, D)** Superimposed structures of 9 and 17 paralogs from the purple Cluster A and grey Cluster B modules, respectively, following the trimming of disordered N- and C-terminal regions. Structural alignment with PyMOL reveals high conservation with mean RMSD values of 3.36 Å ± 1.35 Å for the purple cluster and 4.91 Å ± 1.01 Å for the grey cluster. Sequence similarity based on BLOSUM62 shows average similarities of 50.62 % ± 7.43 % and 51.77 % ± 7.60 %, respectively. Known Carboxypeptidases clustered with the putative novel proteases are P10619, Q9HB40, Q9H3G5. Novel ones in the light-yellow cluster: B are Q8WTS1, Q5VYY2, Q8TB40, P07098, P07099, Q5VXJ0, P38571, Q5W064, Q5VXI9, Q5EB52, Q92597, Q8IUS5, Q9H6B9, Q9BV23. **A noteworthy finding is that all of these have highly similar shapes of their structures**.

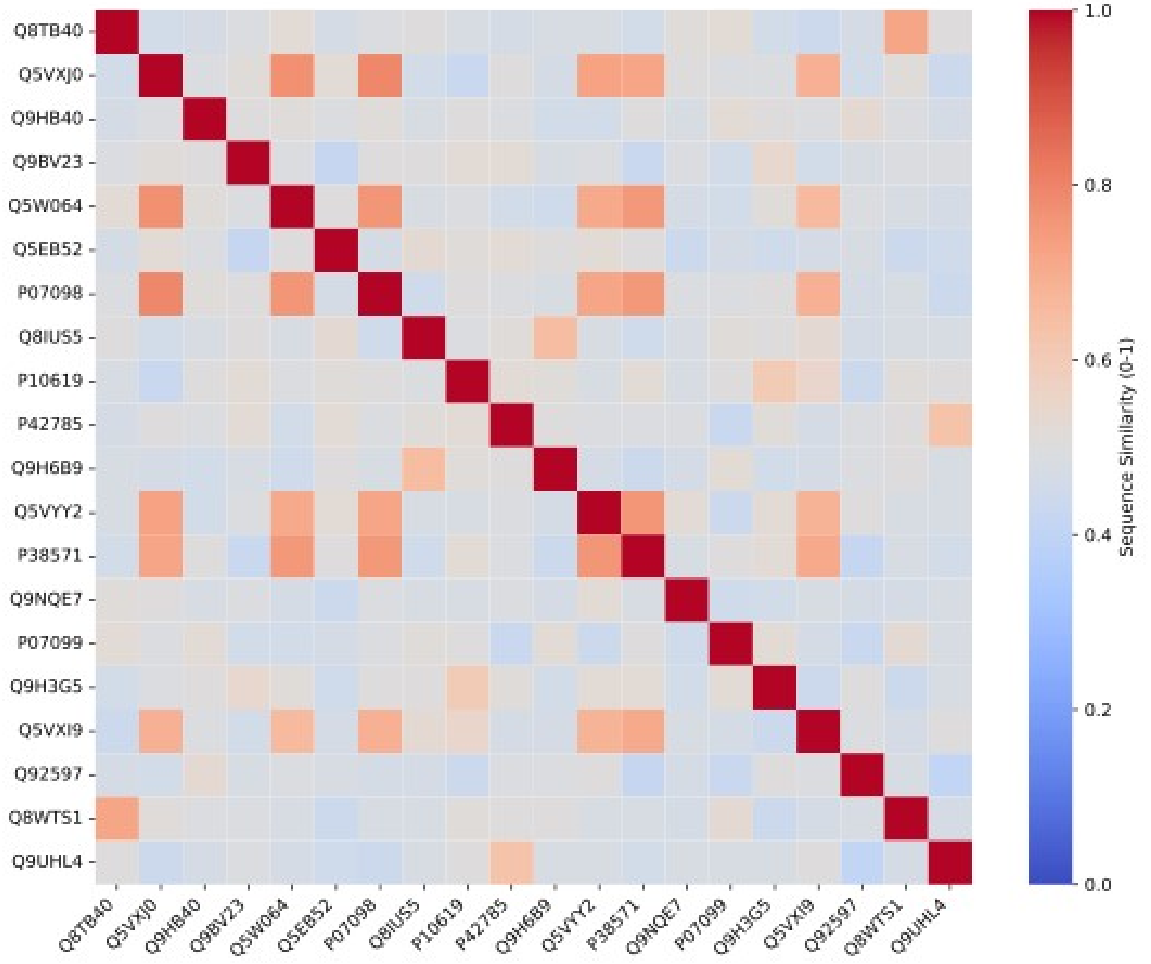


**Figure S3. Heatmap showing sequence similarity of reference and putative novel human serine carboxypeptidases.** The sequence similarity based on BLOSUM62 shows an average similarity of 51.77 % ± 7.60 %

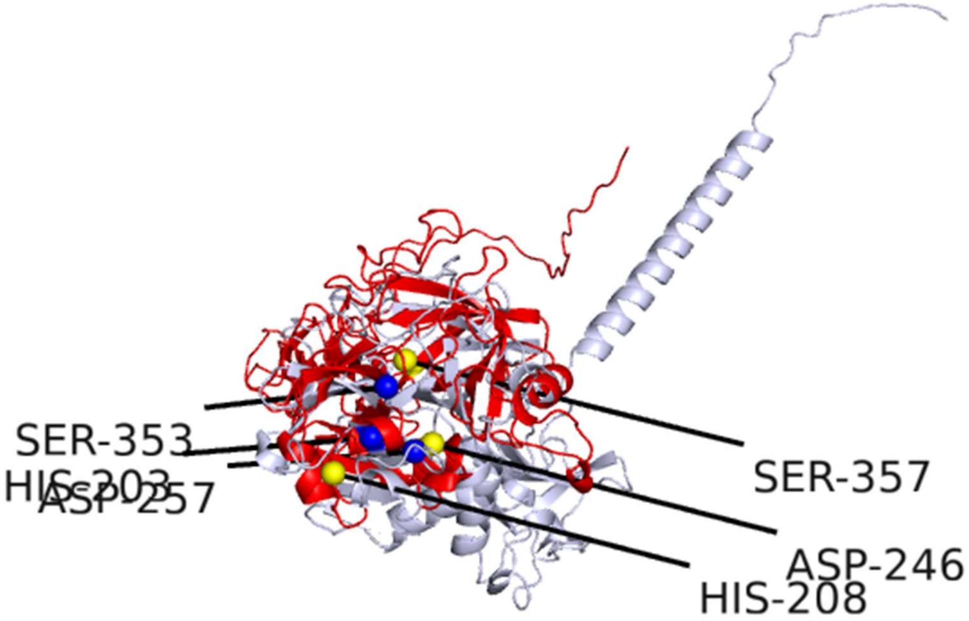


**Figure S4**. Alphafold generated human chymotrypsin-like serine proteases with disordered regions embedded. Reference- P05981 (grey), Target- P00738 (red), RMSD 5.80 Å. Predicted active site residues highlighted in yellow spheres for the target protein.

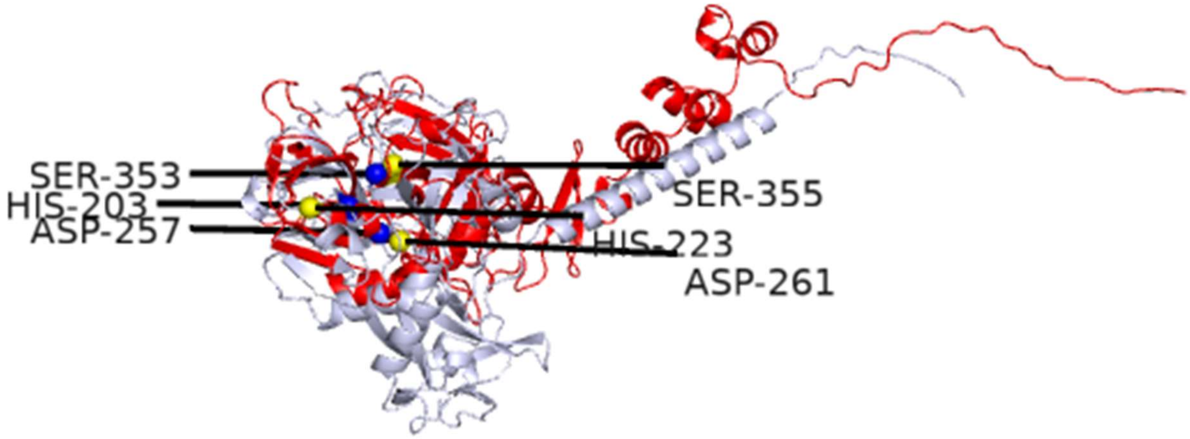


**Figure S5**. Alphafold generated human chymotrypsin-like serine proteases with disordered regions embedded. Reference – P05981 (grey), Target - P22891 (red), RMSD- 4.98 Å. Predicted active site residues highlighted in yellow spheres for the target protein.

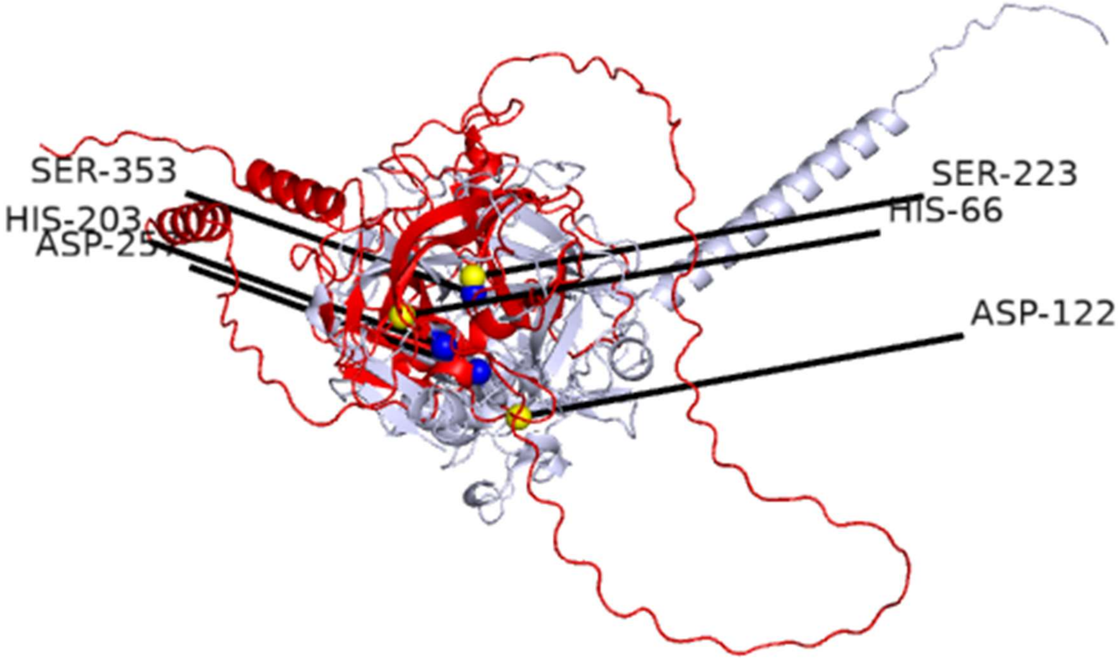


**Figure S6**. Alphafold generated human chymotrypsin-like serine proteases with disordered regions embedded. Reference- P05981 (grey), Target- Q6PEW0 (red), RMSD- 6.88 Å. Predicted active site residues highlighted in yellow spheres for the target protein.

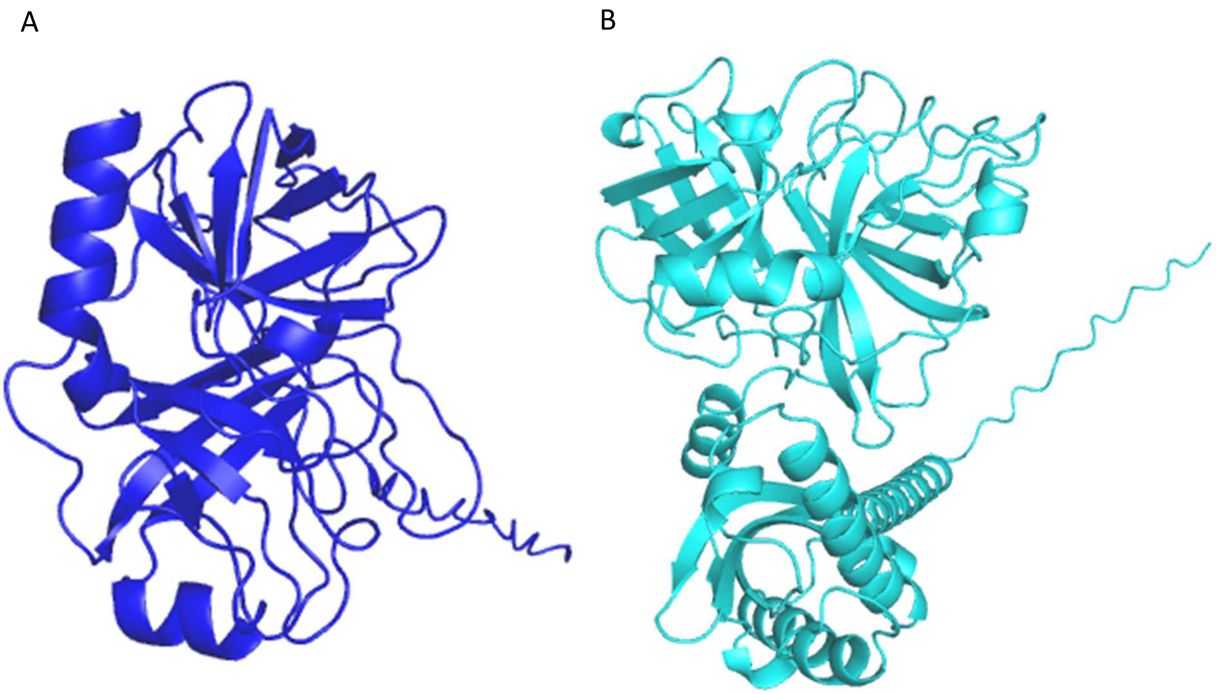


**Figure S7. Size difference in two major human chymotrypsin-like serine protease clusters.** The colors correspond to the clusters in Fig. S1 A. Structure of protein P07477 from blue cluster in Fig. S1 with average sequence length of 269 and B. Protein Q9UL52 from light blue cluster with average sequence length of 395.

**Table S1. 114 of 132 human protease structures clustered into four major clusters in Fig. S1. Blue highlighted are chymotrypsin-like serine proteases and green highlighted are serine carboxypeptidases.** Clusters with members < 5, are excluded for sanity purpose. Novel discoveries are highlighted in yellow.

| Cluster color | #new/#known | Members |
|---|---|---|
| Light blue | 0/36 | A1L453, P04070, Q7RTY5, Q9UL52, Q9H3S3, Q8NF86, Q9NRR2, P40313, Q9BQR3, Q6ZMR5, P08217, P08218, Q6ZWK6, Q99895, P57727, P22891, Q16651, Q6UWB4, P05981, Q86T26, P08709, O60235, Q6PEW0, P00749, P00742, P00738, P00740, Q9NRS4, Q9Y6M0, Q86WS5, P10323, Q9GZN4, O15393, Q9BYE2, Q9NZP8, Q14520 |
| Blue | 0/49 | A4D1T9, P07478, P07477, Q9UNI1, Q9Y5K2, P08861, P20231, P24158, O60259, Q7RTY3, Q6GPI1, P49862, Q9BZJ3, Q9UKR3, Q9P0G3, O43240, P49863, A8MTI9, P17538, Q6UWY2, P00739, A6NIE9, Q9Y337, Q9UKQ9, Q8NHM4, P51124, P12544, P20160, P23946, Q9H2R5, P10144, P09093, P20718, P08246, P07288, P00746, Q92876, P08311, Q9UKR0, Q9UBX7, Q8IYP2, P06870, |

| | | |
|---|---|---|
| | | Q15661, Q9UI38, P35030, P20151, Q7Z5A4, Q7RTY9, A0A1B0GVH4 |
| Light green | 14/6 | P10619, Q5VXJ0, P07098, Q5W064, Q9UHL4, Q5VYY2, P38571, Q8WTS1, Q8TB40, Q9NQE7, Q5VXI9, Q9H3G5, P07099, Q9HB40, P42785, Q8IUS5, Q5EB52, Q9H6B9, Q9BV23, Q92597 |
| Green | 0/9 | P13798, Q86TI2, Q8N608, Q4J6C6, P48147, P27487, P42658, Q12884, Q6V1X1 |

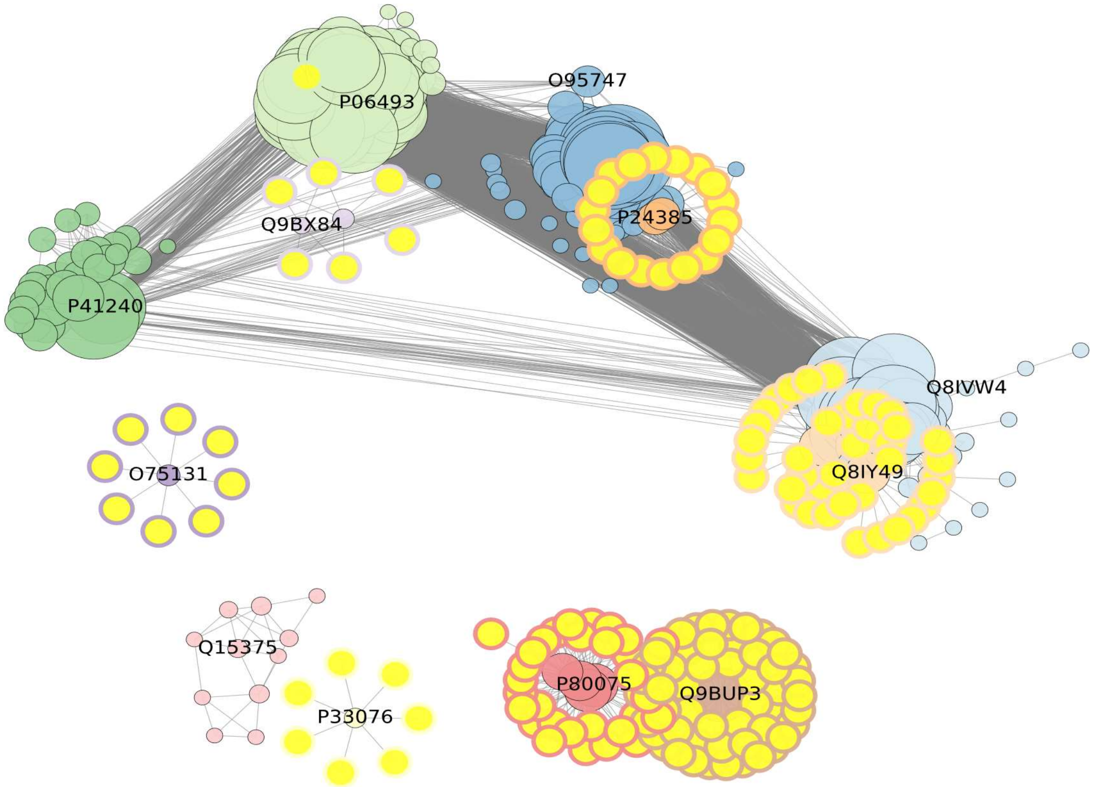


**Figure S8. Network representation of 417 structurally similar (TM-score ≥ 0.5 for both directions) human kinase paralogs showing their complex structural relationships.** 582 proteins with TM-scores ≥ 0.5 bidirectionally clustered by using the Louvain community detection algorithm, for clusters larger than 7 are shown. 163 newly identified kinases are highlighted in yellow with the different corresponding cluster boundary colors are clustered together with the known kinases. Apparently, most of the putative novel kinases are clustered separately, suggesting some entirely new types of kinases.

**Table S2. 417 human kinases distributed in 12 clusters; 163 novel ones highlighted in yellow, green highlighted ones are likely to be NAD biniders based on Uniprot annotation of** Q9BUP3 **as NAD binder**

| Cluster color | # new/# known | Members |
|---|---|---|
| Light green | 1/101 | Q13554, Q13555, P06493, P24941, Q00535, Q00526, P11802, Q00534, P50613, Q15131, P19784, Q8IZL9, Q8N165, Q96PN8, Q59FN2, Q9BXA6, Q6SA08, P36507, Q6P2M8, Q02750, O75716, Q16816, Q9P1W9, Q9BUB5, O94768, Q5MAI5, Q86V86, P15735, P48729, Q9UIK4, P17612, P0C264, Q8NEV1, Q8N752, P0C263, P68400, Q9UQB9, P11309, P52564, Q9C0K7, Q8TDX7, P22612, Q92519, Q8WU08, P22694, Q96QS6, Q7RTN6, P46734, O00506, |

| | | |
|---|---|---|
| | | Q9P289, Q9Y6E0, P51955, Q9NY57, O43293, Q8NCB2, Q8IW41, Q8IY84, Q16644, Q9HBH9, P49137, Q9UEE5, Q15831, Q8N2I9, Q14012, P11801, Q52WX2, Q8IU85, Q9BXA7, Q96PF2, Q6A1A2, P41279, Q96RU7, P78368, Q9HCP0, P51817, Q16566, Q9Y6M4, Q9HC98, Q96GD4, Q8TAS1, P48730, P49674, Q6J9G0, Q9H5K3, Q86YV6, Q96KB5, P00540, O14733, P45985, O43930, O14965, Q96RU8, Q13163, Q96S44, Q96NX5, Q86Y07, Q13557, Q9UQM7, Q8IV63, Q8TDR2, Q99986, Q9H1R3 |
| Green | 0/32 | P42680, P42685, P06239, P41240, Q9Y572, O43353, Q16671, P27037, Q13705, P37023, P37173, Q8NER5, Q04771, P36897, O00238, P36894, P36896, Q06187, Q13882, P07948, P42681, P08631, P12931, P51451, P07947, P09769, P42679, P06241, Q9H3Y6, Q08881, Q13418, Q8NB16 |
| Dark purple | 8/1 | O75131, Q99829, Q8IYJ1, O95741, Q96A23, Q96FN4, Q86YQ8, Q9HCH3, Q9UBL6 |
| Pink | 0/11 | P54764, Q15375, P29320, P29322, P54760, P54753, Q5JZY3, P54756, O15197, Q9UF33, P29323 |
| Yellow | 7/1 | P33076, Q9HC29, Q86W25, Q86W28, Q9NX02, P59046, Q7RTR0, Q96MN2 |
| Red | 36 | P80075, P80098, Q99616, P51671, P13500, Q9Y258, P10147, P55774, P16619, P13236, Q16627, P22362, P13501, Q92583, P10145, Q8NHW4, P78556, P48061, Q99731, P02778, Q16663, O00626, O43927, P55773, Q9Y4X3, P19875, P42830, P09341, P19876, P80162, P02776, O00175, O15467, P47992, P10720, Q9UBD3 |
| Brown | 54 | Q9BUP3, P30043, Q9NZL9, P09417, Q9NUI1, Q9BY49, Q7Z4W1, Q9BPX1, O75828, P26439, P14060, P15428, P35270, O75911, Q8NBN7, Q99714, Q8IZV5, Q8TC12, Q96LJ7, Q8N5I4, Q92781, Q9BUT1, O75452, Q8N3Y7, Q7Z5P4, P16152, Q3SXM5, Q9Y394, P0CG22, A0PJE2, O14756, P56937, Q14376, Q9H2F3, O95455, A6NNS2, Q6IAN0, Q8N4T8, Q92506, Q9HBL8, Q9NRG7, Q6UWP2, Q9NYR8, P37058, Q8IZJ6, Q06136, Q8NBQ5, Q9BTZ2, P28845, Q6PKH6, Q53GQ0, Q96NR8, Q9HBH5, Q13630 |
| Light orange | 40 | Q8IY49, Q15546, Q8N4S7, Q6ZVX9, Q86WK9, Q6TCH7, Q9NUN7, Q9NXK6, Q14656, Q8TEZ7, Q5QJU3, Q6TCH4, Q8TDN7, Q8N661, Q8N2M4, A6NI61, Q9NYV8, P59534, Q9NYW3, Q9NYW2, Q7RTR8, Q8NH60, P59551, P04201, P46089, Q96LB2, Q9Y2T6, P59536, Q8NGN6, Q86SM5, O00270, Q969V1, A6NC51, Q96FM1, Q96LA9, Q8NH56, Q96R54, Q8NG92, Q86V24, Q96A54 |
| Light blue | 44 | O14757, Q8IVW4, Q9UPZ9, P20794, Q16659, P31152, Q6PHR2, P50750, P28482, P53778, Q00532, Q16539, P27361, O15264, Q15759, Q96Q40, O94921, P49841, Q07002, Q00536, P53779, P49336, P49840, P45983, Q9UPE1, P45984, Q92772, Q8TD08, Q9BWU1, Q9UQ07, Q8NE63, Q9NR20, O43781, Q92630, Q00537, Q13627, Q9Y463, Q96SB4, P78362, Q9UBE8, Q9HAZ1, P49759, P49760, P49761 |
| Orange | 20 | P24385, P30279, P30281, P51959, P24863, P51946, Q16589, Q8N815, Q8IV13, Q14094, Q8N1B3, O96020, P24864, Q9H8S5, Q5T5M9, P20248, P14635, O95067, P22674, Q6ZMN8 |
| Blue | 53 | P54646, Q13131, P51956, O95747, Q05513, P41743, O75914, |

| | | |
|---|---|---|
| | | Q04759, Q9NQU5, Q9NWZ3, Q05655, Q13153, O96013, Q13177, Q02156, P24723, Q13188, Q13043, Q9UEW8, Q9BYT3, Q96RR4, Q86UX6, Q8N5S9, O00141, Q99640, P23443, Q9UBS0, Q9Y2H1, Q9HBY8, Q15208, Q96BR1, Q96LW2, O15530, Q8WTQ7, Q15835, P32298, P43250, P34947, P35626, P25098, Q9Y243, P31751, P31749, Q13976, Q13237, P17252, P05771, P05129, Q09013, Q15349, P51812, Q15418, Q9UK32 |
| Grey | 8 | Q9BX84, Q96QT4, Q7Z4N2, Q9HCF6, O94759, Q9NZQ8, Q8TD43, Q7Z2W7 |

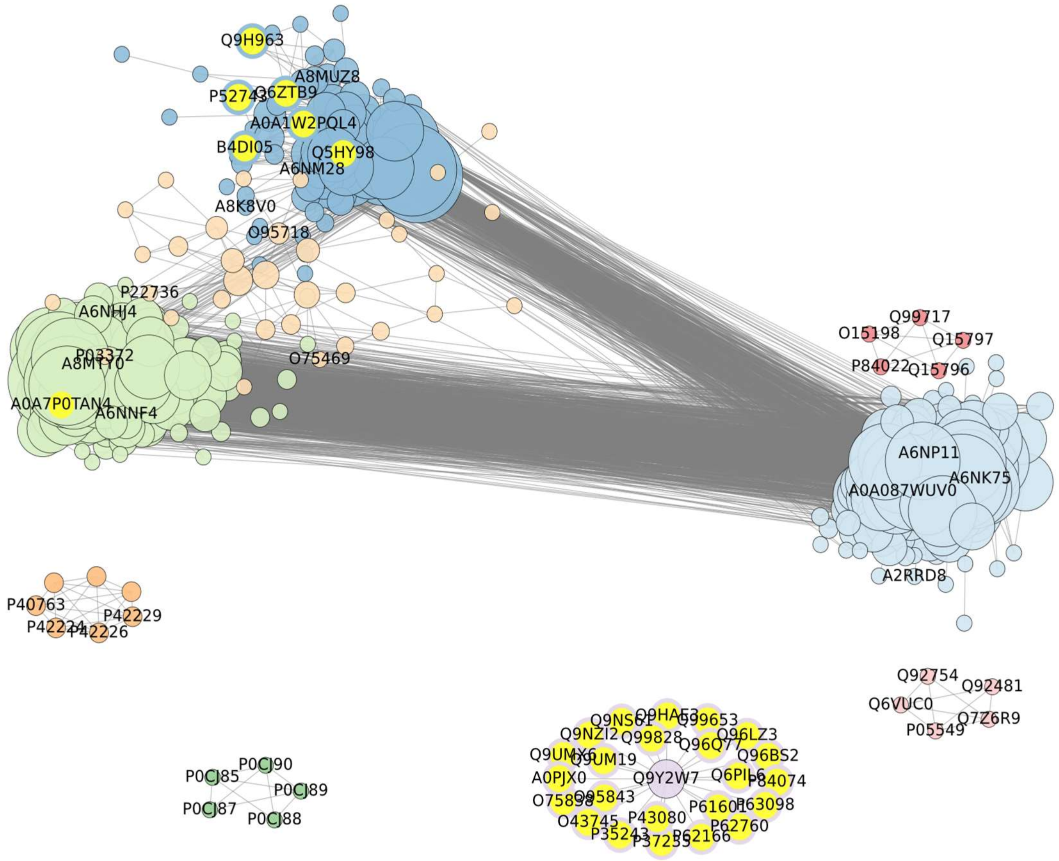


**Figure S9**. **Network representation of 417 structurally similar (TM-score ≥ 0.5 for both directions) human DNA-binding transcription factors and its paralogs.** 30 putative novel TFs highlighted in yellow with the different cluster boundary colors are clustered together with the known TFs.1373 reference proteins (from PANGo) and their 5,061 unique paralogs from integrated homology search after structure-match validation gave 548 proteins. Removing small clusters (size < 5) distributed the remaining 417 nodes into 9 major clusters (members list in **supplementalTable S3**).

| **Table S3. DNA-binding human transcription factor clusters showing the 30 putative novel candidates highlighted in yellow** | | |
|---|---|---|
| Cluster color | Size | Members |
| Light blue | 133 | A0A087WUV0, A2RRD8, A6NK75, A6NP11, A8MUV8, B4DU55, B4DXR9, C9JN71, O75123, O75346, O75373, O95780, P0CG31, P0DPD5, P13682, P15622, P17019, P17024, P17030, P17031, P17032, P21506, P51522, P51786, P52737, P52738, Q03924, Q03936, Q09FC8, Q12901, Q13398, Q14584, Q14585, Q14592, Q14593, Q147U1, Q14929, Q15935, Q15937, Q16587, Q2M3W8, Q2VY69, Q3SXZ3, Q494X3, Q49AA0, Q5SXM1, Q5TEC3, Q5VIY5, Q68DI1, Q68DY1, Q68DY9, Q68EA5, Q6NX45, Q6P1L6, Q6P280, Q6P9A1, Q6PDB4, Q6PF04, Q6ZMS4, Q6ZMV8, Q6ZMW2, Q6ZMY9, Q6ZNH5, Q7L945, Q7Z3I7, Q7Z7K2, Q86UD4, Q86XU0, Q86Y25, Q8IW36, Q8IYI8, Q8IYN0, Q8IZ26, Q8N141, Q8N2I2, Q8N587, Q8N883, Q8N9F8, Q8N9K5, Q8NA42, Q8NB42, Q8NDP4, Q8NEK5, Q8NEM1, Q8NF99, Q8TA94, Q8TAF7, Q8TAU3, Q8TB69, Q8TBZ5, Q8TBZ8, Q8TC21, Q8TD23, Q8WV37, Q96CS4, Q96H40, Q96HQ0, Q96JC4, Q96N38, Q96N58, Q96ND8, Q96NG8, Q96NI8, Q96NJ3, Q96NJ6, Q96NL3, Q96PE6, Q96RE9, Q96SQ5, Q9BR84, Q9BS31, Q9BWM5, Q9C0F3, Q9GZX5, Q9H5H4, Q9H8G1, Q9HBT8, Q9HCL3, Q9NQX1, Q9NQX6, Q9NQZ8, Q9NSJ1, Q9P255, Q9UC06, Q9UC07, Q9UJL9, Q9UK11, Q9UK12, Q9ULM2, Q9Y2G7, Q9Y2P0, Q9Y2Q1, Q9Y6Q3 |
| Blue | 80 | A0A1W2PQL4, A6NM28, A8K8V0, A8MUZ8, A8MWA4, B4DI05, B4DX44, B7ZLF3, P0CB33, P0CH99, P0CI00, P10073, P10075, P17023, P17026, P17036, P17041, P51504, P52741, P52743, P52744, P58317, Q0D2J5, Q13106, Q13360, Q15072, Q15928, Q15940, Q15973, Q16600, Q3KNS6, Q3SY52, Q3ZCT1, Q49A33, Q5HY98, Q5JVG8, Q5T5D7, Q5VV52, Q6P2D0, Q6PK81, Q6U7Q0, Q6V9R5, Q6ZN57, Q6ZN79, Q6ZNG0, Q6ZS27, Q6ZTB9, Q7L3S4, Q7Z398, Q86XF7, Q8IVC4, Q8IVP9, Q8IYX0, Q8N3J9, Q8N782, Q8N8L2, Q8N8Y5, Q8N988, Q8N9Z0, Q8NCK3, Q969W8, Q96BR6, Q96CX3, Q96H86, Q96I27, Q96K75, Q96N20, Q96NG5, Q96SR6, Q9BS34, Q9BSG1, Q9BSK1, Q9H7X3, Q9H963, Q9NR11, Q9NSD4, Q9NXT0, Q9P0T4, Q9UIE0, Q9UJN7 |
| Light green | 124 | A0A7P0TAN4, A6NHJ4, A6NNF4, A8MTY0, A8MXY4, B7Z6K7, O43296, O75290, O75437, O94892, P0CJ79, P0DKX0, P10072, P15621, P16415, P17014, P17017, P17020, P17021, P17025, P17027, P17035, P17038, P17039, P35789, P51508, P51523, P51814, P52736, P52740, P52742, Q02386, Q03923, Q03938, Q06730, Q06732, Q08AN1, Q08ER8, Q0VGE8, Q14586, Q14587, Q14588, Q14590, Q2M218, Q32M78, Q3KP31, Q3MIS6, Q3ZCX4, Q4V348, Q52M93, Q5CZA5, Q5JNZ3, Q5JVG2, Q5MCW4, Q5TYW1, Q6ECI4, Q6IV72, Q6NX49, Q6P3V2, |

| | | |
|---|---|---|
| | | Q6PG37, Q6ZN06, Q6ZN08, Q6ZN19, Q6ZNA1, Q6ZNG1, Q6ZR52, Q76KX8, Q7Z340, Q7Z3V5, Q7Z7L9, Q86T29, Q86UE3, Q86V71, Q86WZ6, Q86XN6, Q86YE8, Q8IYB9, Q8N184, Q8N4W9, Q8N7K0, Q8N7M2, Q8N7Q3, Q8N823, Q8N8J6, Q8N972, Q8NB50, Q8NDQ6, Q8NEP9, Q8NHY6, Q8TAQ5, Q8TF20, Q8TF32, Q8TF45, Q8WXB4, Q96GE5, Q96IR2, Q96K58, Q96MU6, Q96N22, Q96SE7, Q96SK3, Q99676, Q9BRH9, Q9BX82, Q9H0M5, Q9H7R0, Q9H7R5, Q9HBT7, Q9HCG1, Q9HCX3, Q9NYT6, Q9NZL3, Q9P0L1, Q9P2J8, Q9UII5, Q9UJW7, Q9UJW8, Q9UK10, Q9UK13, Q9Y2A4, Q9Y2P7, Q9Y3M9, Q9Y473, Q9Y6R6 |
| Green | 5 | P0CJ85, P0CJ87, P0CJ88, P0CJ89, P0CJ90 |
| Light orange | 33 | P22736, O75469, O95718, P03372, P10276, P10588, P10589, P10826, P10827, P10828, P11473, P11474, P13631, P19793, P24468, P28702, P35398, P37231, P41235, P48443, P51449, P55055, P62508, Q03181, Q07869, Q13133, Q14541, Q14994, Q92570, Q92731, Q92753, Q9Y466, Q9Y5X4 |
| Light pink | 5 | Q6VUC0, P05549, Q7Z6R9, Q92481, Q92754 |
| Pink | 5 | O15198, P84022, Q15796, Q15797, Q99717 |
| Orange | 8 | P40763, P42224, P42226, P42229, P51692, P52630, Q14765 |
| Purple | 24 | Q9Y2W7, A0PJX0, O43745, O75838, O95843, P35243, P37235, P43080, P61601, P62166, P62760, P63098, P84074, Q6PIL6, Q96BS2, Q96LZ3, Q96Q77, Q99653, Q99828, Q9HAE3, Q9NS61, Q9NZI2, Q9UM19, Q9UMX6 |

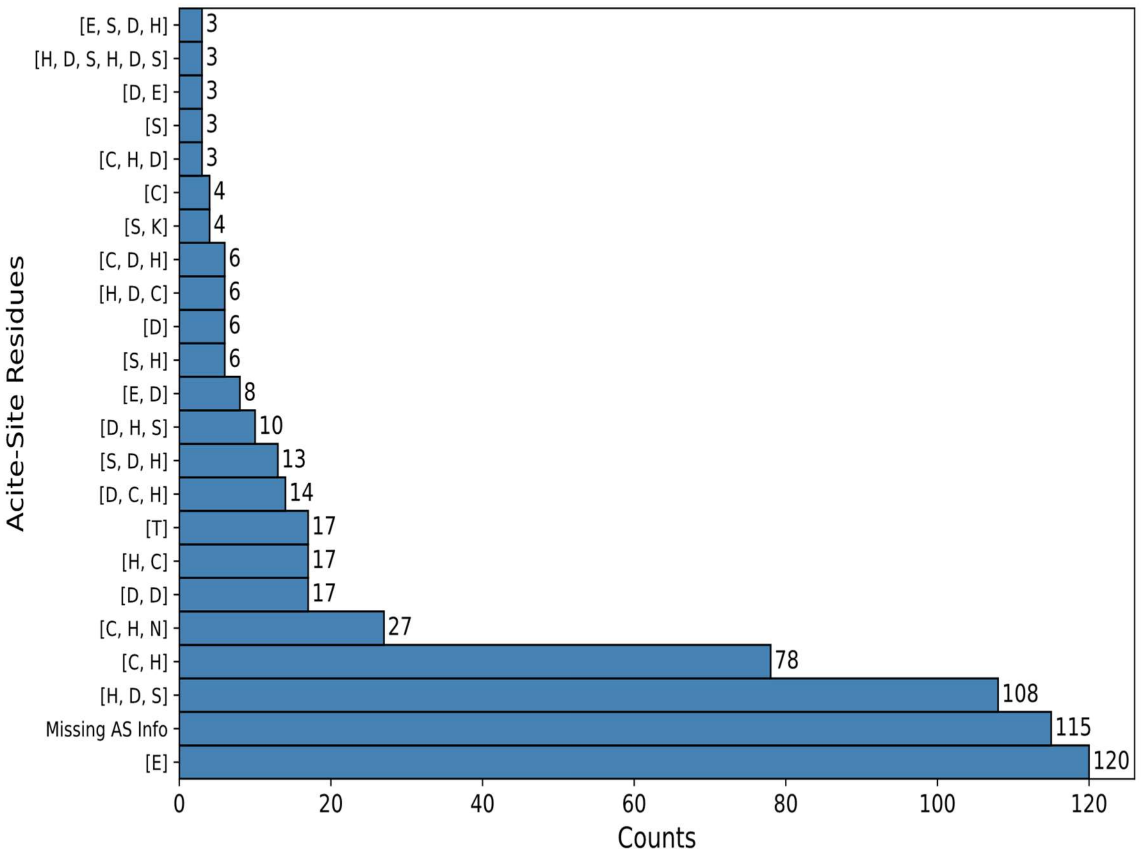


**Figure S10. The 618 human proteases annotated in QuickGO distributed according to their active site residues taken from UniProt.** Active site types with more than 2 proteins only are shown here.

| **Table S5. Summary of Relevant Serine Protease Literature Refer to Table 1 for names and other information** |  |
|---|---|
| **Protein(s)** | **Comment** |
| pseudoproteases | Some of these new paralogs are identified as serine pseudoproteases. One useful is “Serine pseudoproteases in physiology and disease” (1). |
| **Q8WTS1** | Protein ABHD5 does not have serine protease annotation but has been experimentally shown that is a serine protease. “ABHD5 acts *in vivo* and *in vitro* as a serine protease cleaving HDAC4. Through the production of an N-terminal polypeptide of HDAC4 (HDAC4-NT), ABHD5 inhibits MEF2-dependent gene expression and thereby controls glucose handling” (2). |
| **Q5EB52** | Serine protease activity has been suggested but not verified. “MEST has the highly conserved catalytic triad, serine 145-histidine 146-aspartate 147, within the conserved sequence motif for lipases and serine proteases (accession number NP_032616); thus, it is possible that MEST has a role in lipid metabolism, even though its specific enzymatic activity has not yet been identified” (3). |
| **P07098 and Q5W064** | These two proteins are lipases. Lipases have the same type of catalytic triad (Ser-His-Asp) used by serine proteases to break down lipids, but they have different substrates as targets in comparison to serine proteases. One of the first papers to solve the crystal structure of a triacylglycerol lipase is called “A serine protease triad forms the catalytic center of a triacylglycerol lipase”. They revealed “a Ser..His..Asp trypsin-like catalytic triad with an active serine buried under a short helical fragment of a long surface loop”(4). |
| **P00738** | This protein is a haptoglobin. Haptoglobin is considered a serine protease homolog |

| | |
|---|---|
| | but have different biological functions (5). Haptoglobin is regarded as a serine pseudoprotease (1). |
| **P20160** | Azurocidin shares structural similarity with serine proteases but lacks the catalytic activity of a true serine protease, which is crucial for its other functions. Azurocidin is a serine pseudoprotease (1). |

**Table S6. Summary of Relevant Kinase Literature Refer to Table S2 for names and other information**

| Protein(s) | Comment |
|---|---|
| **Q96RU7** | Tribbles homolog 3 (SKIP3 or TRIB3) is a pseudo kinase, meaning it has a structure similar to a kinase but lacks actual kinase activity and the kinase-activation domain (1). |
| **Q99829, Q8IYJ1, O95741, Q96A23, Q96FN4, Q86YQ8, Q9HCH3, Q9UBL6** | These are copines and appear to be identified as kinases because one copine (III) was found to have significant copine activity. (2) |
| **Q9HC29, Q86W25, Q86W28, Q9NX02, P59046, Q7RTR0, Q96MN2** | The proteins do not appear to be kinases but have LRR domain other kinases have |

| | |
|---|---|
| | (specifically the one pre-existing entry in the group). (3) |
| **P80098, Q99616, P51671, Q9Y258, P55774, P16619, P13236, Q16627, P22362, Q92583, P10145, Q8NHW4, P78556, P48061, Q99731, P02778, Q16663, O00626, O43927, P55773, Q9Y4X3, P19875, P42830, P09341, P19876, P80162, P02776, O00175, O15467, P10720, Q9UBD3** | The one preexisting entry was a chemokine as were the above, and the following article was labeled in Uniprot as having "protein kinase activity" but the article itself did not directly note chemokines to have any, as opposed to being in a pathway involving a kinase. (4) |
| **P30043, Q9NZL9, P09417, Q9NUI1, Q9BY49, Q7Z4W1, Q9BPX1, O75828, P26439, P14060, P15428, P35270, O75911, Q8NBN7, Q99714, Q8IZV5, Q8TC12, Q96LJ7, Q8N5I4, Q92781, Q9BUT1, O75452, Q8N3Y7, Q7Z5P4, P16152, Q9Y394, P0CG22, A0PJE2, O14756, P56937, Q9H2F3, A6NNS2, Q6IAN0, Q8N4T8, Q92506, Q9HBL8, Q9NRG7, Q6UWP2, Q9NYR8, P37058, Q8IZJ6, Q06136, Q8NBQ5, Q9BTZ2, P28845, Q6PKH6, Q53GQ0, Q96NR8, Q9HBH5, Q13630,** | Proteins in this class bind (though may be inactive or predicted) NADP/NAD (including but not restricted to adenosyltransferases, dehydrogenases and reductase) The reason there was matching was that the one pre-existing entry in this group was incorrectly noted to have kinase activity, but later discovered to bind NADPH and not ATP. (5) |
| **Q14376** | This protein is an epimerase operating on UDH (Rossman-like fold). (6) |
| **O95455** | This protein is a dehydratase operating on dTDP. (7) |
| **Q8N4S7, Q6ZVX9, Q86WK9, Q6TCH7, Q9NXK6, Q8TEZ7, Q6TCH4, Q86V24, Q96A54** | The common factor here for these are PAQR, with PAQR10 and PAQR11. The following article noted kinases, but not that the receptors themselves had kinase activity.(8) |
| **Q9NUN7, Q14656, Q5QJU3, Q8TDN7, Q8N661, Q8N2M4, A6NI61, A6NC51, Q96FM1** | These is related to PAQR receptors in that both are in the CREST superfamily (note Q14656 was determined using PROST to be ceramidase based on Drosophila melanogaster 3.14E-13 E-value with respect to alkaline ceramidase, Q8N661 to be ceramidase based on Arabidopsis thaliana 2.14E-02 E-value with respect to alkaline ceramidase, Q8N2M4 to be ceramidase based on Arabidopsis thaliana 5.49 E-5 E-value with respect to alkaline ceramidase, A6NI61 to be ceramidase based on Homo sapiens 8.39 E-23 |

| | |
|---|---|
| | E-value with respect to alkaline ceramidase 1, A6NC51 to be ceramidase based on Drosophila melanogaster 1,53E-3 E-value with respect to alkaline ceramidase, Q96FM1 to be ceramidase based on Mus musculus 2.08E-02 E-value with respect to alkaline ceramidase 3. (9)<br><br>Not sure about how taste receptors, olfactory receptors, angiotensin receptor and other miscellaneous in above category fit in (are in same category with PAQR10 and PAQR11. |
| **P30279, P51959, P24863, P51946, Q16589, Q8N815, Q8IV13, Q14094, Q8N1B3, O96020, P24864, Q9H8S5, Q5T5M9, P20248, P14635, O95067, P22674, Q6ZMN8** | Pre-existing entries labeled as kinases but papers cited indicates part of complex involving kinase and not kinase itself. (10,11) |
| **Q7Z4N2, Q9HCF6, O94759, Q9NZQ8, Q8TD43, Q7Z2W7** | The two pre-existing proteins in this entry identified with kinase domain, but other proteins in this family (TRPM) may not have this domain. (12) |

| Table S7. Summary of Relevant Transcription Factor Literature Refer to Table S3 for names and other information |  |
|---|---|
| **Protein** | **Comment** |
| **A0A1W2PQL4, Q5HY98** | **Zn fingers** |
| **P52743, Q5HY98, Q6ZTB9, Q9H963** | Annotated as putative zinc fingers, may be involved in transcriptional regulation. |
| **B4DI05** | UniProt mentions that this protein is highly similar to Zinc finger protein 263. |

**Louvain Community Clustering based on TM-align score**

Due to complex relation between many proteins in terms of similarity, we tried to separate them into different clusters based on the similarity networks defined by the TM-scores using Louvain Algorithm. Here, each protein is represented as node and two nodes are connected by undirected edge if the TM-score is >= 0.5 (for both reference and target) forming network of communities. One of the ways to detect community is by modularity of the community that defines the strength of network. High modularity means more connections between the nodes within a community but sparse connections between nodes of different communities. This method maximizes the modularity of the module or the community. The gain in modularity by moving an isolated node i into a community *C* can be computed by the following equation:

$$\Delta \mathrm{G} = \left[\frac{\Sigma \mathrm{in} \ + \ 2\mathrm{ki,in}}{2m} - \left(\frac{\Sigma \mathrm{tot} \ + \mathrm{ki2m}}{2m}\right)^2\right] - \left[\frac{\Sigma \mathrm{in}}{2m} - \left(\frac{\Sigma \mathrm{tot}}{2m}\right)^2 - \left(\frac{\mathrm{ki}}{2m}\right)^2\right], \quad (1)$$

Where $\Sigma in$ is the sum of the weights of the links inside C, $\Sigma tot$ is the sum of the weights of the links to node in *C*, $k_i$ is the sum of the weights of the links incident to node I, $k_{i, in}$ is the sum of the weights of the links from I to node in *C* and m is the sum of the weights of all the links in the network.
